# Application of Unmanned Aircraft Systems (UASs) for Disease Assessment and High Throughput Field Phenotyping of Plant Breeding Trials

**DOI:** 10.1101/2025.06.16.659708

**Authors:** Tadesse Anberbir, Felix Bankole, Girma Mamo, Gerald Blasch, Alemu Dabi

## Abstract

Conventional plant phenotyping relies on visual scoring and manual measurements, which are labor-intensive, time-consuming, and prone to human error. To address these limitations, Unmanned Aircraft Systems (UASs) are increasingly being applied in breeding trials to capture various phenotypic traits. High Throughput Phenotyping (HTP) offers enhanced speed, accuracy, and efficiency, while potentially reducing costs in plant breeding programs.

This study explores UAS-based phenotyping in wheat breeding trials with the aim to integrate HTP platforms across breeding pipelines. UAS images were acquired using a Parrot Bluegrass drone equipped with a sequoia multispectral sensor, processed via Agisoft Metashape (open-source) and Pix4D mapper (licensed), and analyzed using PlotPhenix (licensed) for Vegetation Indices (VIs) and plot segmentation.

Comparisons between UAS-based and ground-based measurements revealed that grain yield is significantly negatively correlated (r = -0.74) with yellow rust disease severity. Multispectral-derived indices, particularly **Red, Red Edge, and NIR bands, showed positive correlations with grain yield (ranging from 0.22 to 0.23), though RGB-generated indices exhibited stronger correlations.

The findings confirm that UAS-generated indices effectively assess yellow rust disease severity and predict grain yield. UAS-based phenotyping enhances efficiency and accuracy in trait collection and disease assessment, facilitating the development of improved wheat varieties and promoting the integration of UAS technologies into breeding programs.

## Introduction

Plant phenotyping refers to the measurement and assessment of physically observable traits of a plant. Traditionally, this process relies on visual scoring and manual measurements, which are labor-intensive, time-consuming, and prone to human error. As a result, the quantitative analysis of plot-level metrics and crop modeling research has been constrained, especially in large breeding trials, where extracting various metrics presents significant challenges.

To address these limitations, Remote Sensing (RS) platforms, such as Unmanned Aircraft Systems (UASs), have recently gained traction in agricultural data collection systems. These platforms enable high-throughput phenotyping (HTP) by capturing diverse biophysical, biochemical, and sanitary traits, which can be used to predict and explain yield outcomes (Guo et al.). Compared to traditional end-of-season trait assessments, RS technologies offer greater precision in breeding and research, particularly for studying genotype responses to new environmental conditions. With HTP methods, breeding programs can expect enhanced speed, accuracy, and efficiency, along with potentially reduced costs. As these technologies continue to advance, they hold immense promise for improving breeding strategies and optimizing agricultural productivity.

High Throughput Phenotyping (HTP) represents a transformative approach in plant breeding, enabling researchers to enhance efficiency and adapt to future agricultural challenges. Despite its benefits, digital phenotyping technologies remain predominantly adopted in developed nations, while their accessibility in developing countries is hindered by technical expertise, resource constraints, and performance evaluation challenges. Integrating HTP technologies is essential for strengthening national crop breeding programs, across the world, as food security and wellness remains a priority.

Research in digital phenotyping technologies have not been fully explored and more awareness among researchers and practitioners are needed. For instance, Gebremariam et al., (2018) explored UAS imagery using machine learning algorithms for detecting stem rust disease in wheat crops, however, a significant gap in research related to UAS applications in HTP for breeding trials existed. Traditional manual phenotyping methods remain the standard, despite their labor-intensive, error-prone, and time-consuming nature. Particularly, wheat disease assessment and severity measurements pose considerable challenges due to the cost, labor demands, and scalability limitations associated with manual evaluations.

This study aims to introduce UAS-based phenotyping to improve high-resolution data capture and analysis for wheat breeding trials. The overarching goal is to accelerate genetic gain by integrating Artificial Intelligence (AI)-assisted HTP platforms and digitization into breeding programs. This transition seeks to replace manual data collection with automated, high-throughput phenotyping solutions by leveraging existing digital platforms.

The research was conducted at Bekoji sub-station, where wheat breeding trials were performed to screen high-yielding, disease-resistant genotypes under natural rust disease infestation. UAS-generated vegetation indices were correlated with ground-based assessments of yellow rust severity and grain yield, highlighting UASs’ potential in crop health and productivity analysis. The results indicate a significant negative correlation (r = -0.74) between grain yield and yellow **rust** severity, whereas multispectral vegetation indices (Red, RedEdge, and NIR) exhibited positive correlations with yield (ranging from 0.22 to 0.23). However, RGB-based indices demonstrated stronger correlations with grain yield.

Overall, this study validates the application of UAS-based phenotyping as an effective tool for trait collection and disease severity assessment, enhancing efficiency and precision in wheat breeding trials. The findings underscore the potential of UAS technology to support breeding programs and facilitate the development of improved wheat varieties.

## Site Survey, Materials and Methods

The field experiment was conducted at Bekoji sub-station, under the Kulumsa Agricultural Research Center (KARC) (See Figure 1). This location was designated for screening wheat genotypes with high yield potential and disease resistance, particularly against natural infestation of rust diseases.

**Figure. 1.**
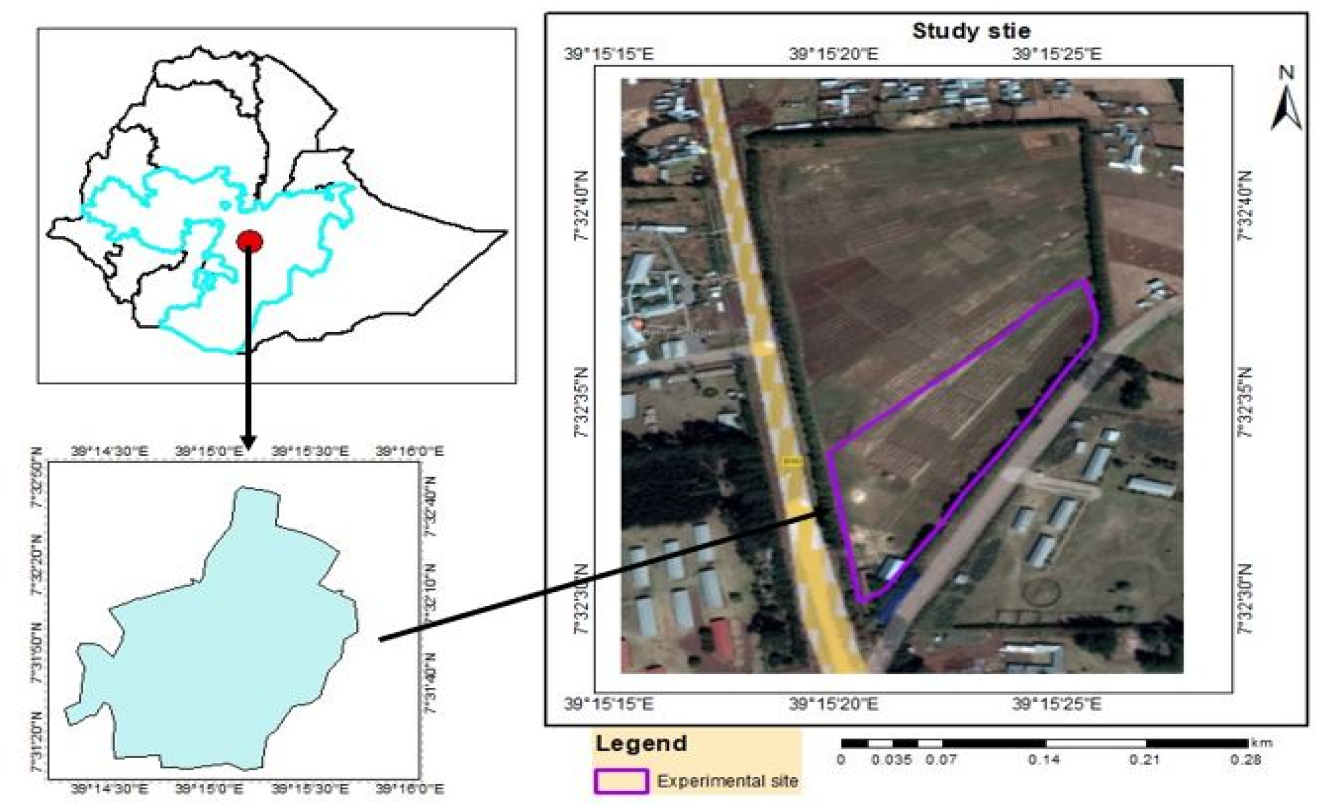
Location of the study area

### Location and Climate

- Geographical Coordinates: 39°15’20.5”E / 7°32’42.0”N
- Altitude: 2,200 meters above sea level
- Region: Bekoji zone, Oromia Region, Central Ethiopia

The site experiences an annual rainfall of 820 mm, following a unimodal distribution pattern. The average air temperature ranges between 10.5°C and 22.8°C, providing a growing period of 120–135 days (Solomon, 2021).

### Disease Prevalence

The climatic conditions in the region are particularly favorable for Yellow Rust (YR) disease occurrence. With YR being endemic in Ethiopia, wheat fields within the KARC area are naturally prone to Yellow Rust infection during the Meher season.

### Experimental Design

The trial was conducted using a randomized complete block design, featuring 80 wheat genotypes. The experiment included four blocks, each with two replicates, resulting in a total of 942 plots. (See Figure 2).

**Figure. 2:**
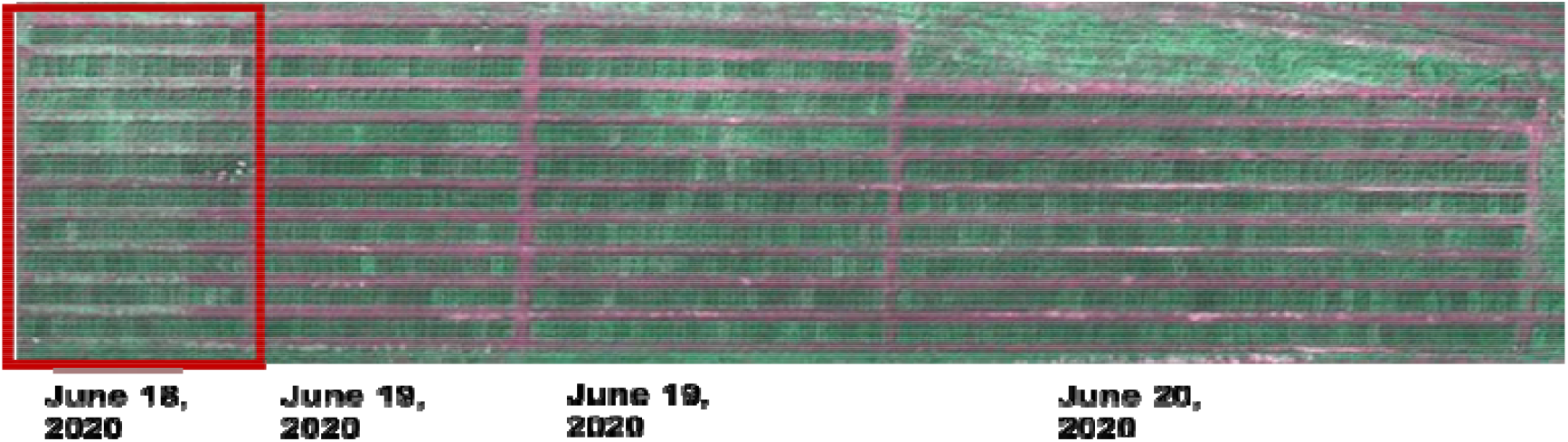
Experimental field layout with four blacks & planting date (Orthomosiac image generated from UAS aerial image)

### Plot Arrangement

- Each block contained ten rows per plot, with dimensions of 1.2 m × 2.5 m, and a 50 cm spacing between plots.
- The trial comprised 80 entries, with one variety per plot and 10 cm spacing between planted rows.
- The first block, consisting of 160 plots, was selected for focused study.

### Management Practices

- Seed Rate: 150 kg ha_ □ ^1^ per plot.
- Fertilizer Application: Applied at a rate of (to be specified).
- Agronomic Practices: Uniform management was implemented across all plots.

### Disease Exposure

- None of the blocks received disease treatments, allowing for natural disease infestation.

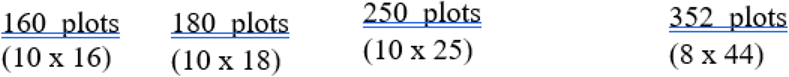

### Data Collection and Analysis

Yellow Rust (YR), a destructive biotic disease, poses a significant threat to wheat crops worldwide, leading to substantial yield losses. Understanding the relationship between yellow rust severity and grain yield is essential for effective disease management and breeding strategies. To achieve this, analyzing vegetation indices (VIs) extracted from multispectral imagery alongside ground truth data offers a valuable indirect approach to predictive analytics.

In this study, field data was collected from wheat sites exhibiting varying degrees of yellow rust severity, where visual disease scoring and grain yield measurements were conducted. Multispectral images were acquired using UAS-mounted satellite-based sensors, capturing spectral bands such as Near Infrared (NIR), Red Edge, Red, Green, and Blue. Preprocessing of imagery including radiometric and geometric corrections, along with orthorectification to ensure accuracy and consistency were required. Vegetation indices such as NDVI and EVI were extracted and analyzed, providing insights into the correlation between spectral data and disease severity. By integrating these indices with observed yellow rust severity, predictive models were developed to assess the impact on grain yield. Validation was conducted using independent test datasets to ensure reliability.

Ultimately, the interpretation of results helped identify optimal indices for disease severity prediction, informing breeding programs and precision agriculture strategies aimed at mitigating the impact of yellow rust. The workflow of image collection and processing is shown in Figure 3.

**Figure. 3.**
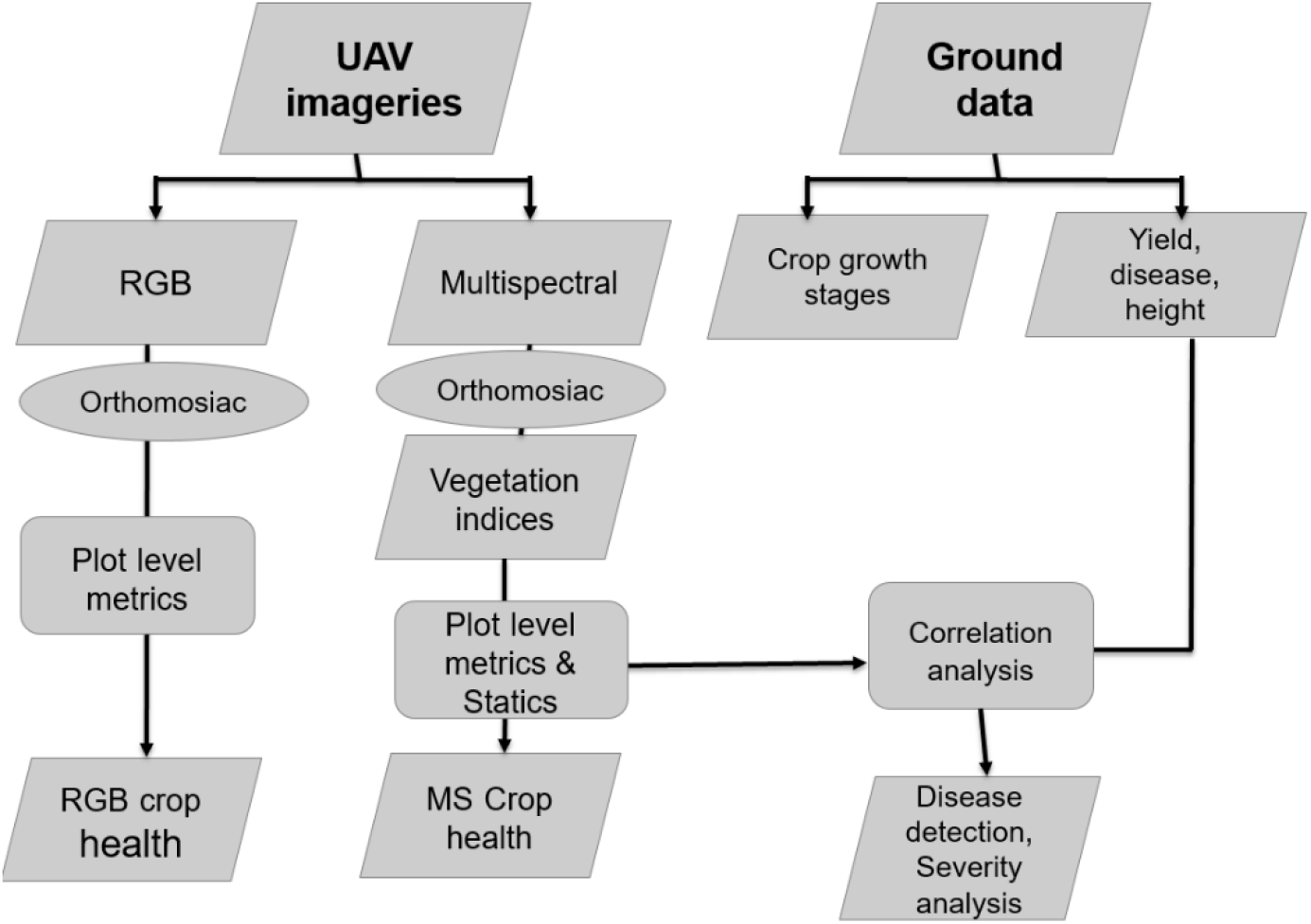
Workflow of image data collection and image processing.

The following highlights steps in data collection and analysis process:

### Ground Data & Disease Scoring

To monitor disease pressure, disease assessments (ground truthing) were conducted throughout the experimental period. Subsequent scoring dates will be determined based on disease development in the plots. Scoring was carried out by researchers, including breeders and pathologists, focusing on disease-related traits such as yellow rust incidence, severity, and response in each plot. After harvesting, yield data were collected (see Table 1).

**Table 1:**
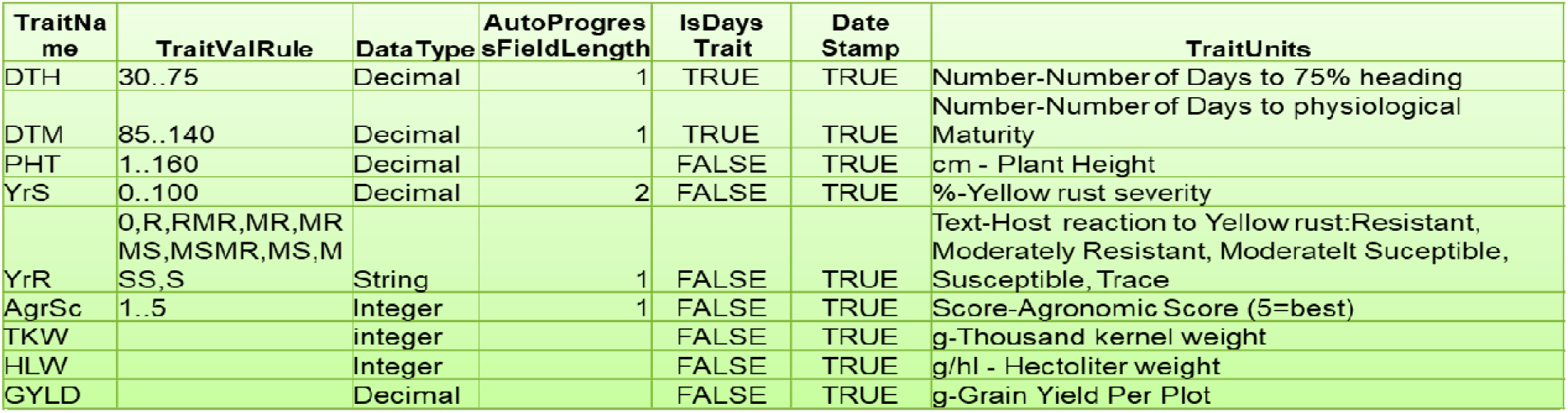
List of major traits.

The critical growth stage was identified under close supervision, considering various growth and yield determinant factors. These included:

- Soil and drainage analysis to assess potential nutritional deficiencies
- Crop health assessments evaluating major disease occurrences such as yellow rust, stem rust, leaf rust, fusarium head blight (FHB), and septoria

For evaluating rust disease severity, standardized rust scoring guidelines were adopted from Prescott et al. (1986), Stubbs et al. (1986), and Roelfs et al. (1992). Visual sub-scores were collected for each plot, with observations taken at four corners and the center. These assessments captured rust severity and the host plant response to each rust disease. The Modified Cobb Scale (Roelfs et al., 1992) was employed to record the percentage of rusted tissue, using a rating scale from 1–100%.

### UAS Aerial Image Acquisition

Multispectral UAS sequoia sensor images were captured in four rounds at different growth stages—tillering, head emergence, flowering, and maturity during the main season. Images were taken on the same day as ground scoring or within one day before or after, except for round 4, where images were acquired with a three-day difference.

Each image was captured with the following parameters:

- Forward overlap: 70%
- Lateral overlap: 80%
- Flight speed: 3-8 m/s
- Flight altitude: 20-30 m above ground

Table 2 provides details on image acquisition, including resolution.

**Table 2:** UAS-images acquisition dates and resolutions.

|  | Round 1 | Round 2 | Round 3 | Round 4 | Round 5 |
| --- | --- | --- | --- | --- | --- |
| **Date** | Aug 13, 2020 | Sep 3, 2020 | Oct 8, 2020 | Oct 13, 2020 | Nov 21, 2020 |
| **Flight height** | 20m | 20m | 20m | 30m | 30m |
| **Resolution**<br>**(cm/px) GSD** | 1.98 | 1.98 | 1.98 | 2.83 | 2.83 |
| **Flight speed** |  |  |  |  |  |
| **Overlaps**<br>**(front/side)** | 80/70 | 80/70 | 80/70 | 80/80 | 80/80 |
| **No. images**<br>**(both RGB & MS**<br>**images)** | 1,467 | 1, 452 | 767 | 732 | 877 |

The flight pattern was designed using Pix4D capture software, ensuring structured data collection. Before each flight, the drone was calibrated using a radiometric calibration panel from the ground control points showing the flight pattern (See Figure 5). To geo-reference aerial images, eight ground control points (GCPs) were strategically placed across the field at the beginning of the season (see Figure 4). The GCPs consisted of 20 cm × 20 cm black-and-white laminated paper, mounted on a 50 cm post for visibility.

**Figure. 4.**
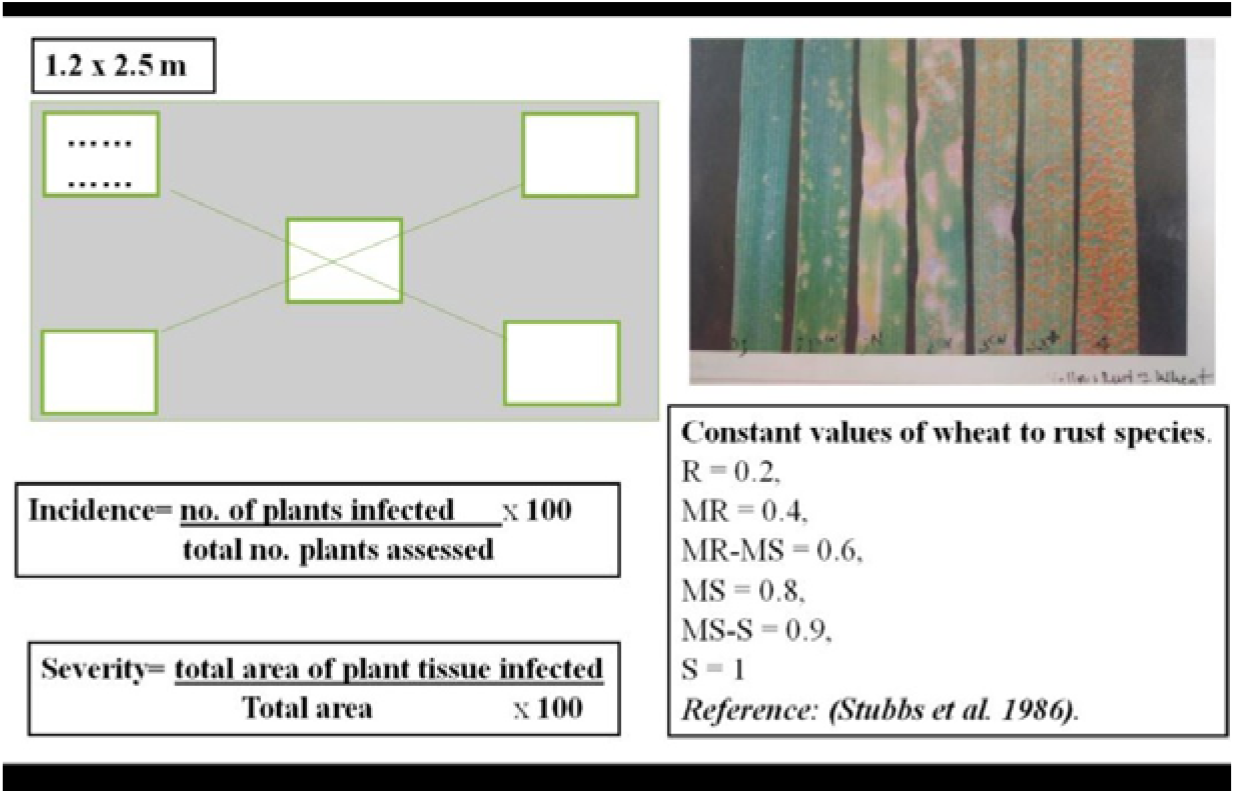
Yellow Rust Disease Incidence and Severity Scoring

**Figure. 5:**
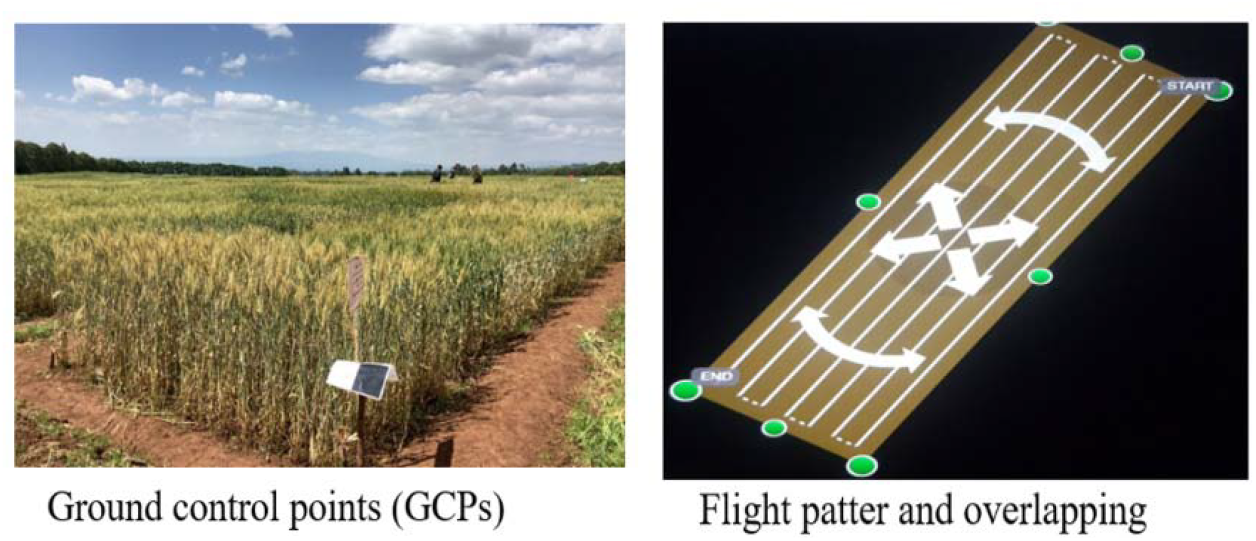
Ground Control Points

The black-and-white contrast ensured easy identification by image processing software. GCP coordinates were measured using a handheld Gemini GPS, ensuring accurate spatial reference for image analysis.

### Image Processing

The parrot sequoia multispectral sensor used in this study captured four discrete spectral bands: Red, Green, Near-Infrared (NIR), and Red Edge. Figure 6 presents mosaicked images of these bands. The captured images were processed using Agisoft Metashape (*open-source, version 1*.*4*.*2*) and Pix4D discovery mapper (*licensed, version 4*.*6*.*4*). Various vegetation indices, including plot segmentation, were generated during processing.

**Figure. 6.**
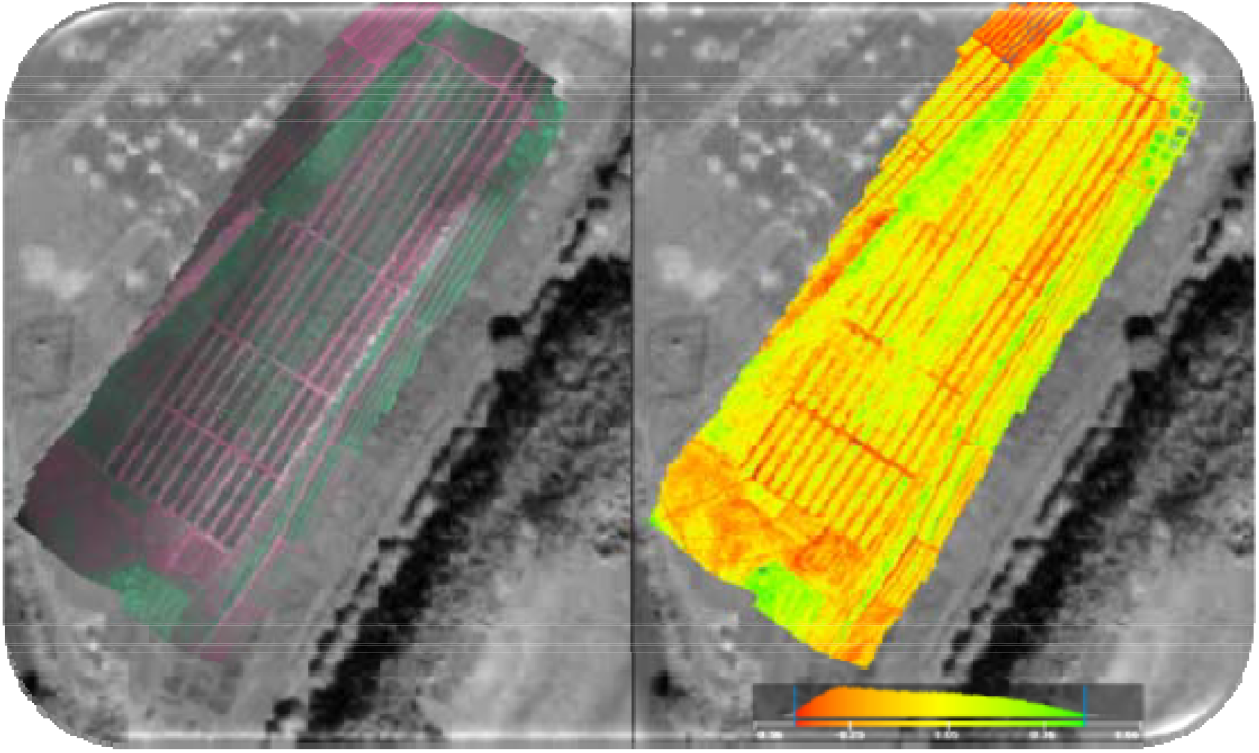
The wheat field images generated from the first round UAS aerial images in: (a) orthomosiac and (b) NDVI at early growth stage

**Figure. 7.**
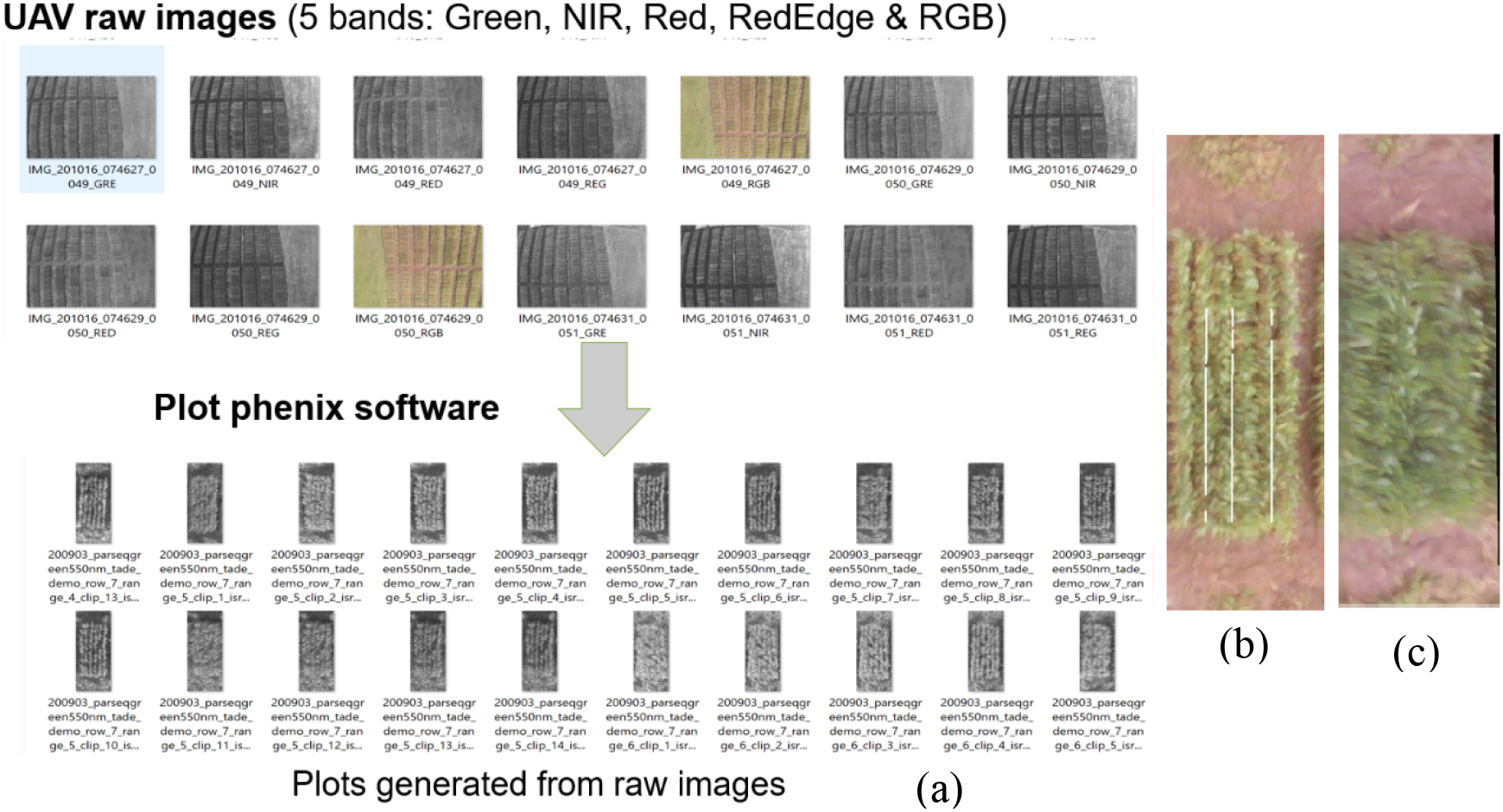
(a) Sample plots generated from UAS images; (b) Sample plot with showing row the six rows and three row length clearly while, (c) shows blurred image due to strong wind

Orthomosaic images were developed using Pix4D mapper, while Plotphenix software was utilized to generate multiple images of each plot taken from different camera perspectives during UAS flights. Figure 6, illustrates the whole experimental field, showcasing both orthomosaic and NDVI images generated from aerial imagery captured at the early growth stage (*August 13, 2020*).

The parrot sequoia multispectral sensor used in the study captured four (4) discrete spectral bands: Red, Green, NIR, and Red Edge. Figure 6 shows mosaicked images of four bands. The images were processed using a photogrammetric software, Agisoft Metashape (open source, version: 1.4.2) and Pix4D discovery mapper (licensed, version 4.6.4) and different vegetation indices were generated, including plot segmentation. Orthomosaic images were generated using Pix4D mapper and Plotphenix software was used to generate multiple images of every plot taken from different camera perspectives during UAS flight. See figure 6 which shows the whole experimental field in which an orthomosiac and NDVI images generated from aerial images taken at the early growth stage (August, 2020).

### Field plot segmentation and vegetation indices extraction

To obtain useful information about each plot in the field, plot-level data needed to be extracted from the orthomosaic image. Individual plot boundaries were defined separately from the images, with an assigned plot ID that identified their genomic type. Plot boundaries were extracted using a grid-based plot extraction method in Plotphenix, a licensed software. Plot-clips, consisting of multiple independent images of the same plot, were extracted to ensure the most accurate calculations of row length under vegetation, percent canopy cover, and reflectance values. Based on these extracted images, vegetation indices such as NDVI and NDRE were calculated. For each plot, eight to nine plot-clips were generated, and the best plot was selected for further analysis.

In addition to these features, Plotphenix also generates average vegetation cover, row length, and various vegetation indices for each plot. Figure 8 presents an average vegetation cover color map generated for the second block, which contains 180 plots. Plot-level data provide wheat breeders with real-time insights into crop performance, enabling them to improve future varieties more precisely and efficiently. Breeders can visualize their field remotely from their office and monitor the performance of each plot using color maps generated for different variables. For example, the plot at (10, 10) has the smallest average vegetation cover, recorded at 65.0. Upon identifying this, the breeder can visit the field to examine the specific plot and investigate potential issues affecting its performance.

**Figure. 8.**
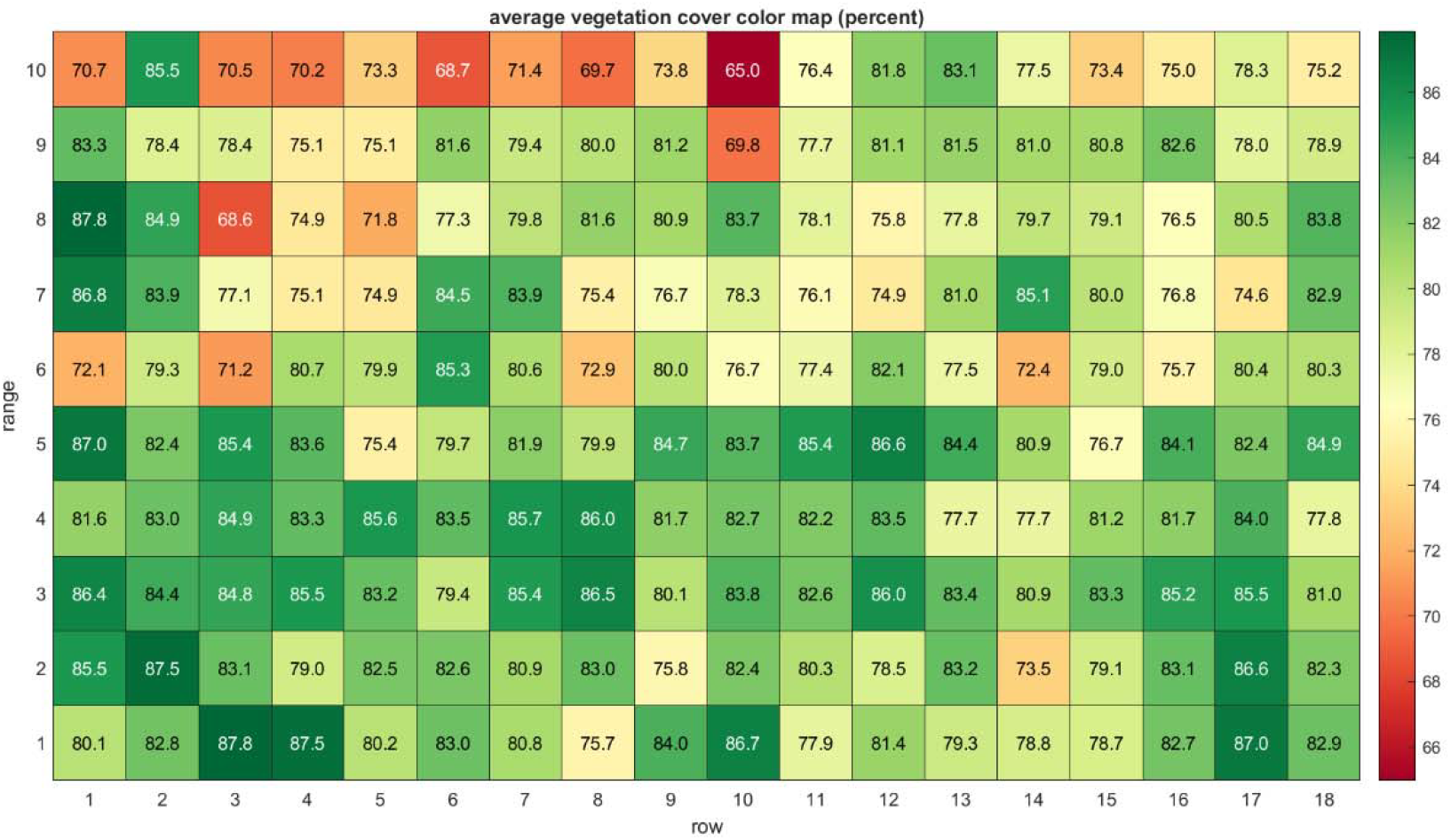
Average vegetation cover in percent for each plot

Moreover, the coefficient of infection was calculated, this is necessary in plant pathology to quantify the severity of disease in relation to its resistance to the disease. Therefore, to obtain a representative rust scoring value per plot, rust severity and host response data were combined in a two-step process.

First, the coefficient of infection (CI) was calculated for each sub-score per scoring date. Then, the resulting CI values were averaged across the plot per scoring date, generating the average coefficient of infection (ACI).

The ACI was calculated following the methods of Saari and Wilcoxson (1974) and Pathan and Park (2006), using the multiplication of disease severity (DS)—a visual estimate of the percentage of leaf area affected by yellow rust—and a constant value representing infection type (IT). The constant values for infection types were assigned as follows:

Where: **R** = 0.2, **MR** = 0.4, **MRMS** = 0.6, **MS** = 0.8, **MSS** = 0.9, **S** = 1

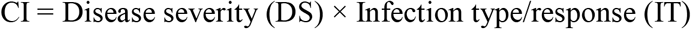

Yellow rust was prioritized over stem rust, as stem rust is rare and largely insignificant under the cold temperatures of study area (Bokoji.).

A statistical analysis was performed using predictive analytics approach to determine the relationship between ground truthing data and Vegetation Indices (VIs), quantifying both the strength and direction of their correlation with grain yield (GYLD). The VIs analyzed in this study included Normalized Difference Vegetation Index (NDVI), Green Normalized Difference Vegetation Index (GNDVI), Modified Soil-Adjusted Vegetation Index (MSAVI), Red Edge Vegetation Index (RDVI), Visible Atmospherically Resistant Index (VARI), Green-Red Vegetation Index (GRVI), Green Leaf Index (GLI), and Normalized Green–Red Difference Index (NGRDI) **(see** Table 3).

**Table 3.** Vegetation Indices calculated from RGB and Multispectral images (adopted from ..)

| Index | Formula | Traits | Relevant Studies |
| --- | --- | --- | --- |
| Green normalized difference vegetation index (GNDVI) | $\frac{(R_{NIR} - R_G)}{(R_{NIR} + R_G)}$ | disease (YR), chlorophyll, LAI, nitrogen, water content | 10,42,43,45 |
| Modified soil adjusted vegetation index (MSAVI) | $0.5[2R_{NIR} + 1 - \sqrt{(2R_{NIR} + 1)^2 - 8(R_{NIR} - R_R)}]$ | disease (YR) | 45 |
| Normalized difference vegetation index (NDVI) | $\frac{(R_{NIR} - R_R)}{(R_{NIR} + R_R)}$ | disease (YR), chlorophyll, LAI, biomass, yield | 10,42–45 |
| Normalized green red difference index (NGRDI) | $\frac{(R_G - R_R)}{(R_G + R_R)}$ | chlorophyll, biomass, water content | 62 |
| Re-normalized difference vegetation index (RDVI) | $\frac{(R_{NIR} - R_R)}{\sqrt{(R_{NIR} + R_R)}}$ | disease (YR) | 10 |
| Green Leaf index (GLI) | $\frac{(R_{2G} - R_b - R_R)}{(R_G + R_R)}$ | | |
| Green-red Vegetation index (GRVI) |  |  |  |
| Visible atmospherically resistant index (VARI<sub>G</sub>) | $\frac{(R_G - R_R)}{(R_G + R_R)}$ | disease (YR) | 10,43,44 |

Note: $R$ : measured reflectance at the original wavelength or wavebands specified by the subscript (G: Green; R: Red; NIR: near-infrared).

For ground truthing data, Plant Height (PHT), Coefficient of Infection Index 1 (CI_1), and Coefficient of Infection Index 2 (CI_2) were considered. The Spearman’s correlation coefficient (**r**) between VIs and ground truthing data was analyzed (Table 4), revealing a significant negative correlation between grain yield and yellow rust disease severity (CI_1 and CI_2) ranging from -0.72 to -0.73. This finding underscores the detrimental effects of yellow rust disease on grain yield, emphasizing the necessity of effective disease management strategies to optimize crop productivity (Figure 9).

**Table 4.**
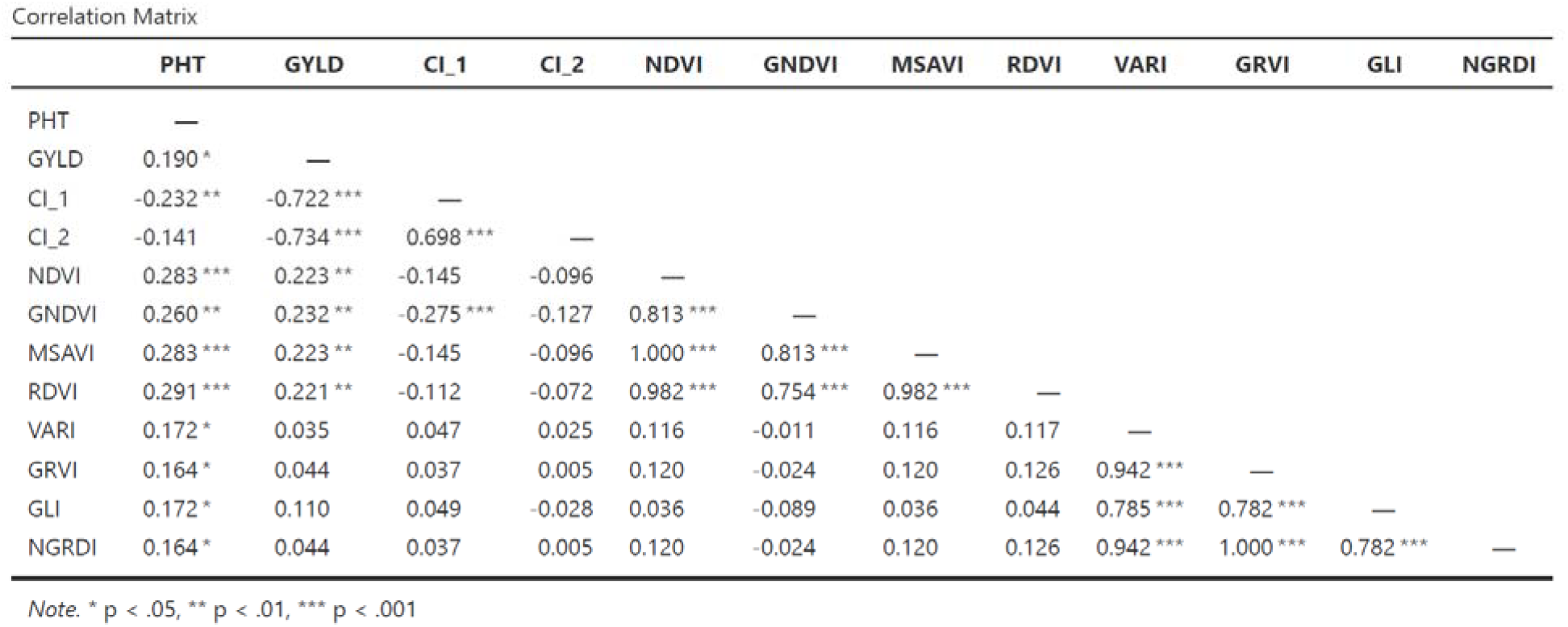
Correlation matrix table between ground data and different vegetation indices.

**Figure. 9.**
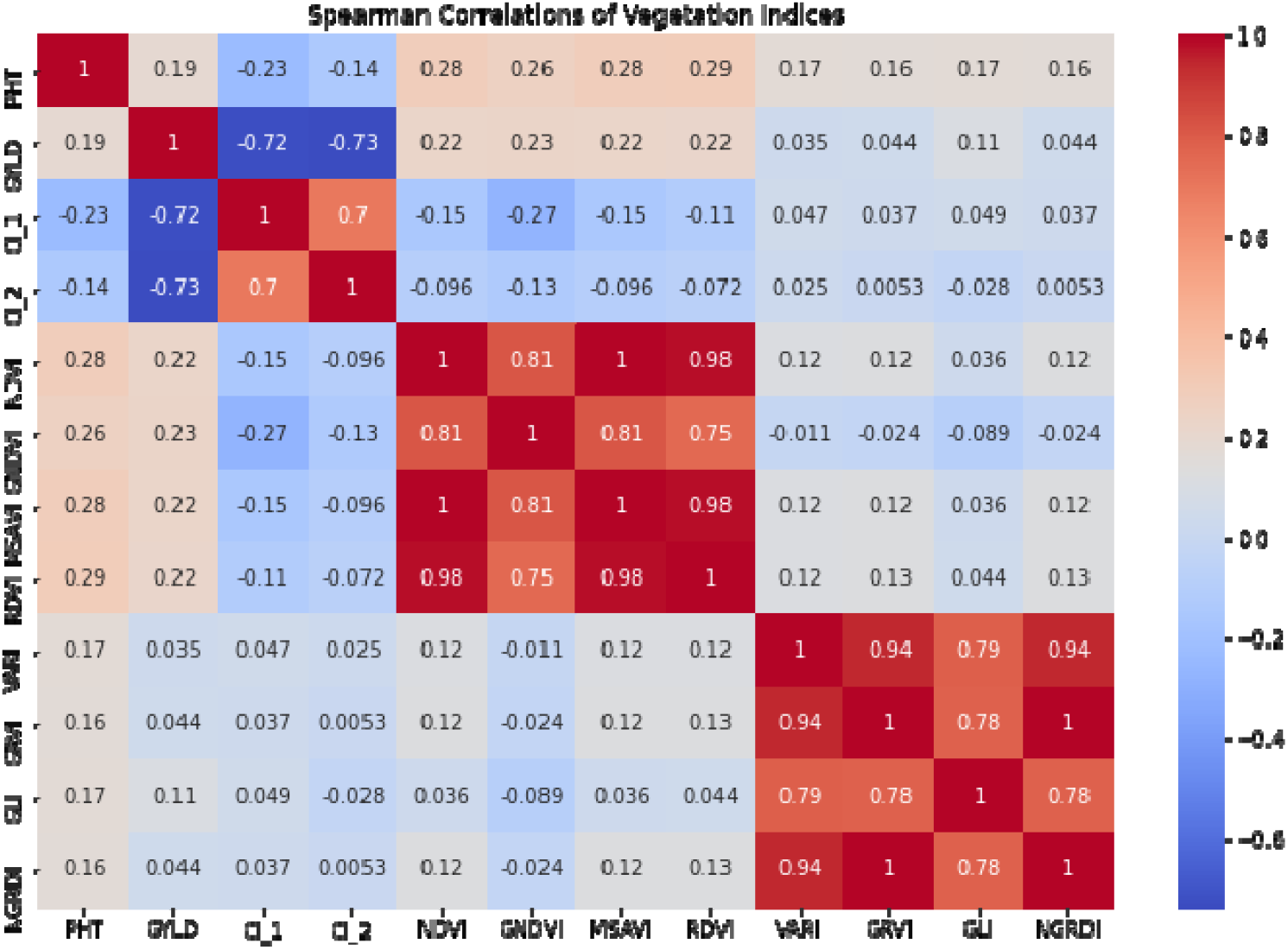
Correlation matrix plot between ground data and different vegetation indices

Additionally, the study found that vegetation indices derived from multispectral images (Red, Red Edge, and NIR) exhibited stronger correlations with GYLD compared to indices generated from RGB images, aligning with previous research findings. Among the indices examined, NDVI, GNDVI, MSAVI, and RDVI showed positive correlations with grain yield, albeit with relatively weak magnitudes (ranging from 0.22 to 0.23). While these indices may hold some predictive value for grain yield, further studies are required to validate these findings (Figure 9).

### Implications for Wheat Breeding and High-Throughput Phenotyping

The positive correlations observed between certain vegetation indices suggest that these indices can provide valuable insights for predicting grain yield. However, caution must be exercised when using them as predictors, as grain yield is influenced by various factors, including soil conditions, weather patterns, and management practices.

Conversely, the strong negative correlation between yellow rust disease (CI_1 and CI_2) and GYLD highlights the significant impact of this disease on crop productivity. This finding underscores the necessity of effective disease management strategies to mitigate yield losses.

These insights hold substantial implications for wheat breeding programs and high-throughput phenotyping. By considering yellow rust severity as a contributing factor to yield loss and utilizing vegetation indices—particularly NDVI—as indirect predictors of grain yield, breeders can make informed decisions to enhance crop productivity.

Additionally, high-throughput phenotyping techniques, such as UAS-based multispectral imaging, offer a non-destructive and efficient method for monitoring crop health and predicting yield potential, enabling precision agriculture advancements.

## Conclusion

In this study, Unmanned Aircraft Systems (UASs) were tested in wheat breeding trials to explore their potential application in the Ethiopian Institute of Agricultural Research (EIAR). The goal was to mainstream high-throughput phenotyping (HTP) platforms and digitization technologies into breeding programs.

Results from UAS-collected data were correlated with manual field measurements, confirming the accuracy and reliability of UAS technology. Our findings demonstrate that UAS-based phenotyping significantly enhances the efficiency and accuracy of trait collection and disease severity assessment in wheat breeding trials, ultimately contributing to the development of improved varieties.

Overall, the experiment confirmed the potential of UAS-based phenotyping for high-throughput data collection in small-scale wheat breeding trials. However, challenges such as data quality limitations, drone-related technical issues, and computational constraints affected the study. Further research is necessary to optimize data collection and analysis protocols and scale up the technology for larger breeding programs.

The application of UASs for high-throughput phenotyping in wheat breeding trials in Ethiopia presents a transformative opportunity to revolutionize the breeding process and enhance the efficiency of variety development. This technology enables breeders to identify superior genotypes more quickly and accurately, facilitating the development of improved wheat varieties that contribute to food security and support smallholder farmers’ livelihoods.

To accelerate the breeding process, crop breeding scientists and pathologists should prioritize the integration of UAS-based technologies. Further research is required to strengthen the relationship between remotely sensed plant phenological traits and biological field measurements, such as plant height, biomass, and yield. Moving forward, we will continue refining phenotyping capabilities to support large-scale experiments and breeding trials, **e**nsuring continuous advancements in precision agriculture and crop improvement.

As part of our future research, we plan to conduct a comparative analysis of wheat high-throughput phenotyping in Western Canada—including Alberta, Manitoba, and Saskatchewan— in comparison with the Ethiopian climate region. This analysis will provide valuable insights into how environmental factors influence wheat growth, disease susceptibility, and yield potential, further enhancing breeding strategies for diverse agricultural conditions.

## Acknowledgments

The authors express their sincere gratitude to the Ethiopian Institute of Agricultural Research (EIAR) for generously funding this project. We also extend our appreciation to the dedicated Kulumsa Agricultural Research Center (KARC) breeding team for their invaluable collaboration in conducting this research. Special thanks to the Plotphenix team (www.plotphenix.com) for their essential support with software and data assistance.

## Reference

Blasch, G., Anberbir, T., Negash, T., et al. (2023). The potential of UAV and very high-resolution satellite imagery for yellow and stem rust detection and phenotyping in Ethiopia. Scientific Reports, 13, 16768. 10.1038/s41598-023-43770-y

Alemu, W., et al. (2016). Effect of environment on wheat varieties, yellow rust resistance, yield, and yield-related traits in South-Eastern Ethiopia. Plan, 4(3).

Hailu, E., et al. (2015). Distribution of stem rust (Puccinia graminis f.sp. riici) races in Ethiopia. Plan, 3(2).

Agricultural Transformation Agency - ATA, Ethiopia. (n.d.). Ethiopian Institute of Agricultural Research. https://www.ata.gov.et

Guo, W., Carroll, M. E., Singh, A., Swetnam, T. L., Merchant, N., Sarkar, S., Singh, A. K., & Ganapathy Subramanian, B. (2021). UAS-based plant phenotyping for research and breeding applications. Plant Phenomics, 2021, Article ID 9840192. 10.34133/2021/9840192

Tadesse, W., Endalamaw, H., Debele, T., Kassa, D., Shiferaw, W., Solomon, T., Negash, T., Geleta, N., Bishaw, Z., & Assefa, S. G. (2022). Wheat production and breeding in Ethiopia: Retrospect and prospects. Crop Breeding Genetics and Genomics.

Jang, G., Kim, J., Yu, J.-K., Kim, H.-J., Kim, Y.-H., Kim, D.-W., Kim, K.-H., Lee, C., & Chung, Y. S. (2020). Review: Cost-effective unmanned aerial vehicle (UAV) platform for field plant breeding applications. Remote Sensing, 12(998). 10.3390/rs12060998

Singh, R. P., Hodson, D. P., Jin, Y., Lagudah, E. S., Ayliffe, M. A., Bhavani, S., et al. (2015). Emergence and spread of new races of wheat stem rust fungus: Continued threat to food security and prospects of genetic control. Phytopathology. 10.1094/PHYTO-01-15-0030-FI

Ganeva, D., Roumenina, E., Dimitrov, P., Gikov, A., Jelev, G., Dragov, R., Bozhanova, V., & Taneva, K. (2022). Phenotypic traits estimation and preliminary yield assessment in different phenophases of wheat breeding experiment based on UAV multispectral images. Remote Sensing, 14(1019). 10.3390/rs14041019

Xie, C., & Yang, C. (2020). A review on plant high-throughput phenotyping traits using UAV-based sensors. Computers and Electronics in Agriculture, 178.

Arias Rojas, F. (2018). Exploring machine learning for disease assessment from high-resolution UAV imagery.

Koc, A., Odilbekov, F., Alamrani, M., et al. (2022). Predicting yellow rust in wheat breeding trials by proximal phenotyping and machine learning. Plant Methods, 18(30).

Haghighattalab, A., González Pérez, L., Mondal, S., et al. (2016). Application of unmanned aerial systems for high-throughput phenotyping of large wheat breeding nurseries. Plant Methods, 12(35). 10.1186/s13007-016-0134-6

Volpato, L., Pinto, F., González-Pérez, L., Thompson, I. G., Borém, A., Reynolds, M., Gérard, B., Molero, G., & Rodrigues, F. A. Jr. (2021). High-throughput field phenotyping for plant height using UAV-based RGB imagery in wheat breeding lines: Feasibility and validation. Frontiers in Plant Science, 12, 591587.

Solomon, T. (2021). Correlation and path coefficient studies on advanced bread wheat lines in Ethiopia. Science Publishing Group, 9, 20. 10.11648/J.CB.20210902.11

Roelfs, A. P., Singh, R. P., & Saari, E. E. (1992). Rust diseases of wheat: Concepts and methods of disease management.

